# Molecular Landscape of the Mouse Adrenal Gland and Adjacent Adipose by Spatial Transcriptomics

**DOI:** 10.1101/2025.01.02.631086

**Authors:** Małgorzata Blatkiewicz, Szymon Hryhorowicz, Marta Szyszka, Joanna Suszyńska-Zajczyk, Andrzej Pławski, Adam Plewiński, Andrea Porzionato, Ludwik K. Malendowicz, Marcin Rucinski

## Abstract

The adrenal glands play a vital role in maintaining homeostasis and managing stress through the production and secretion of steroid hormones. Despite extensive research, the molecular mechanisms underlying adrenal zonation and cellular differentiation remain poorly understood. By employing spatial transcriptomics, this study has mapped the adult CD1 IGS mouse adrenal gland, thereby identifying unique genetic markers of zonal differentiation and dynamic cellular interactions. Five cellular clusters, corresponding to the cortex and medulla compartments, were identified, along with two adipose tissue clusters (brown and white). These findings confirm the centripetal differentiation model, highlighting the gradual transition of cell populations from the capsule through cortical zones. Through ligand-receptor interaction analysis, a complex regulatory network governing inter- and intra-zone communication was identified, thereby emphasising the adrenal gland’s central role in integrating endocrine and neuroendocrine signals, particularly in response to stress. This comprehensive spatial transcriptomic map of the adult mouse adrenal gland provides original insights into adrenal biology and constitutes a valuable resource for future research.

## Introduction

The adrenal glands are a key component of the endocrine system, and they maintain homeostasis within the body, regulating diverse physiological processes and enabling the body to respond effectively to stressors (Guasti *et al*, 2011; Nishimoto *et al*, 2010). The adrenals are surrounded by rich deposits of adipose tissue and are subdivided into the outer cortex and inner medulla. The human adrenal cortex derived from mesoderm is composed of three overlapping concentric zones: zona glomerulosa (ZG), zona fasciculata (ZF), and zona reticularis (ZR) (Miller & Auchus, 2011; Neville & O’Hare, 1985; Nussdorfer, 1986; Yates *et al*, 2013). Diverse anatomical and functional areas within the adrenal cortex synthesize specific steroid hormones in depending on endocrine and paracrine signals. The ZG produces mineralocorticoids, such as aldosterone, which are responsible for regulating blood pressure by managing electrolyte balance(Miller & Auchus, 2011). On the other hand, the ZF synthetizes glucocorticoids, like cortisol, which influence metabolism and stress responses. The innermost ZR, produces a small amount of androgens, which are precursors of sex hormones (Bird, 2012). Adrenal medulla, originated from the neuroectoderm (neural crests) produces and secretes catecholamines responsible for maintaining the body’s homeostasis under conditions of stress.

The adrenal glands of humans and mice exhibit notable differences, particularly in the composition of their cortical zones. In humans, the fetal zone (FZ) and the X-zone (ZX), present in mice, are transient structures within the adrenal cortex(Goel *et al*, 2014; Huang & Kang, 2019). The fetal zone, which is dominant during the fetal period, undergoes rapid regression within a few weeks after birth and at a lower intensity this process lasts more than two years(Huang & Kang, 2019; Malendowicz, 2010). On the other hand, the ZX in the mouse adrenal gland emerges postnatally and its regression depends on the sex (Beuschlein *et al*, 2012; Grabek *et al*, 2019; Ruggiero *et al*, 2021). In females, the ZX gradually disappears over a few months, but it disappears rapidly during pregnancy while in males, it disappears before the 40th day of life(Huang & Kang, 2019). Therefore, the regression of the ZX in mice demonstrates a distinct sexual dimorphism, indicating the potential involvement of hormonal factors (androgens) in this process(Gannon *et al*, 2019). Additionally, evidence suggests that the involution of the ZX may also vary depending on the mouse strain. Despite these differences, mice serve as an excellent research model for the analysis of adrenal function, exhibiting a broad translational potential, including in immunological contexts.

It has been confirmed that the adrenocortical cells undergo continuous renewal from a progenitor pool where cells from the periphery of the ZG migrate centripetally into the inner zones of the adrenal cortex(Lerario *et al*, 2017; Lyraki & Schedl, 2021; Pihlajoki *et al*, 2015). This displacement and differentiation to the inner steroidogenic zones are essential for the repopulation and maintenance of adrenal function due to stress, apoptosis, or developmental changes. Adrenal cells adapt to the production of appropriate steroid hormones by undergoing a specific molecular and metabolic transformation. Conventional molecular biology methods do not fully visualize this process. The presence of this phenomenon has been confirmed via lineage tracing analyses (Freedman *et al*, 2013; King *et al*, 2009; Laufer *et al*, 2012; Pihlajoki *et al*., 2015), but the mechanisms underlying the centripetal differentiation model remain incompletely understood. The continuous transition of the adrenal cells has only been confirmed in a human adrenal to show the spatial relationships between the cells (Rui *et al*, 2023).

Spatial transcriptomics represents a transformative approach in molecular biology, enabling researchers to study gene expression patterns within their native tissue environment(Stahl *et al*, 2016; Wang *et al*, 2020). This innovative technology integrates spatial information with transcriptomic data to provide a comprehensive understanding of cellular heterogeneity, tissue architecture and the functional roles of different cell types in both physiological and pathological contexts(Stahl *et al*., 2016; Wang *et al*., 2020). The visualisation and quantification of gene expression *in situ* have profound implications for the understanding of complex biological systems, particularly in organ-specific studies.

The chosen research model (adrenal glands of adult male mice) allows us to determine the pattern of gene expression in the adrenal glands in which the transient X-zone occurs. It should be noted that we did not find similar studies on mouse adrenal glands (with Visium CytAssist technology) addressing this issue in the available literature. Our studies have identified molecular pathways specific to individual zones of the adrenal gland of adult male mice. They also revealed characteristic signalling pathways within the ZX that are absent, or their expression is very low in the other components of the gland studied. The results of our study extend the knowledge of adrenal biology in which the ‘transient structure (ZX)’ occurs. They may also lead to the search for new factors characterising adrenal biology under both physiological and pathological conditions. As the field continues to evolve, the integration of spatial transcriptomic data with traditional histological and biochemical approaches will undoubtedly enhance our understanding of the complex biology of the adrenal gland.

## Results

### Biomarkers of adrenal components

The spatial gene expression analysis by Visium CytAssist technology was performed on the histological sections of the adrenal glands of male mice (aged 10 weeks, CD1 IGS). Four histological sections of the adrenal glands were analysed; however, for clarity, we present only one section through the adrenal. According to the Space Ranger count summary metrics, the tissue had 992 spots, with an average of 49011 reads per spot and a median of 5904 genes per spot, and a total of 19250 genes were detected.

Using the Uniform Manifold Approximation and Projection (UMAP) data reduction method implemented in the Seurat library, we identified seven distinct cell clusters, shown in different colours on **Figure 1**. Based on the identification of markers for each cluster and their spatial localisation within the adrenal histological slides, we determined the specific cell types associated with each cluster. We identified a clusters characteristic of brown adipose tissue (BAT), white adipose tissue (WAT) and connective tissue (CT – adrenal connective tissue capsule). In UMAP plot, these clusters were located the close to each other, indicating relative similarity. Taking into account the cut-off criteria of average logFC>1 and adj.*p* < 0.05, a total of 3460 markers were obtained, including 342 for ZG, 291 for ZF, 261 for ZX, 1775 for medulla, 258 for CT, 154 for WAT, 379 for BAT. The full marker list is provided in an Excel file as supplementary material (Supplementary File 1). The ten most highly expressed genes characteristic of each cluster are visualised as a heat map, with two representative genes shown on the histological slide. Among them, we identified *Smpx* (small muscle protein, X-linked) and *Ecrg4* (esophageal cancer-related gene 4) for ZG; *Susd3* (sushi domain containing 3) and *Cited1* (CBP/p300-interacting transactivator with Glu/Asp-rich carboxy-terminal domain 1) for ZF; *Sbsn* (suprabasin) and *Ly6m* (lymphocyte antigen 6 complex, locus M) for ZX; *Jph4* (junctophilin 4), *Gnb3* (guanine nucleotide-binding protein subunit beta-3), *Mast1* (microtubule associated serine/threonine kinase 1), and *Tnr* (tenascin R) for the medulla; *Inmt* (indolethylamine N-methyltransferase) and *Mmp9* (matrix metallopeptidase 9) for CT; *Cidec* (cell death-inducing DFFA-like effector c) and *Lep* (leptin) for WAT; and *Aspg* (asparaginase) and *Ucp1* (uncoupling protein 1) for BAT. These spatial expression patterns reinforce the zonation and functional compartmentalisation observed within the adrenal gland, thereby elucidating how gene expression profiles correspond to specific anatomical locations and are involved in adrenal function. We provide also interactive web application to visualize the spatial transcriptomic profile of the mouse adrenal gland. This application allows users to explore the spatial expression profiles of all genes within the dataset, overlaying gene expression data onto high-resolution images of adrenal gland tissue sections.

**Figure 1.**
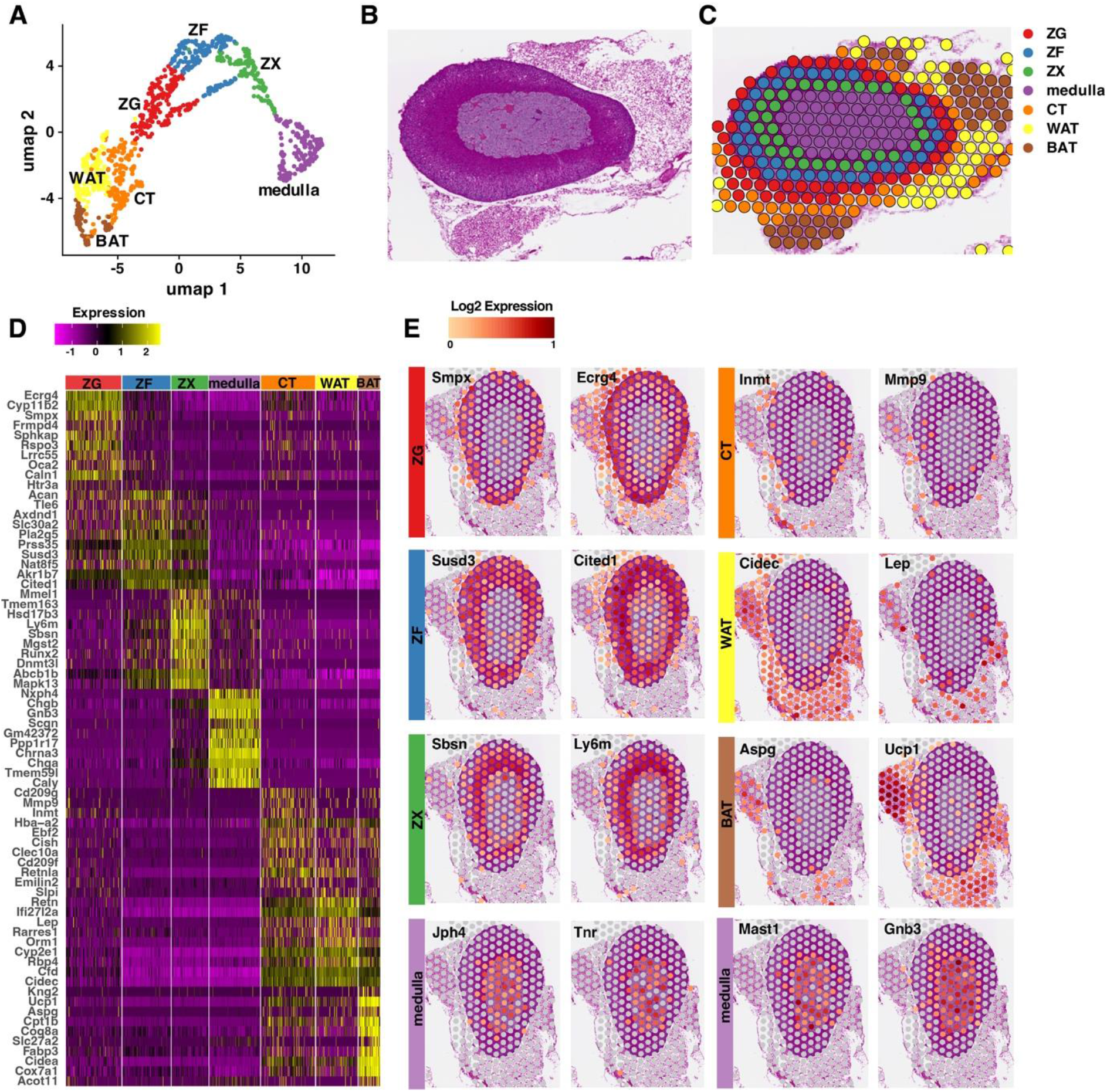
Transcriptomic landscape of the adrenal gland of an adult male mouse. A) Uniform manifold approximation and projection (UMAP) visualisation of adrenal gland spots with manual annotation based on molecular marker prediction for each cluster. BAT – brown adipose tissue, WAT – white adipose tissue, CT – connective tissue, ZG – zona glomerulosa, ZF – zona fasciculata, ZX – X-zone. B) Adrenal glands stained with hematoxylin and eosin (H&E). C) Clusters from the UMAP representation were depicted as color-coded dots on the representative spatial transcriptomic adrenal slide. D) Reconstruction of distinct cell populations within mouse adrenal slices based on UMAP cluster generation. Heat map showing the top 10 expressed (z-score normalised) genes in each obtained cluster. E) Selected representative histological slides with overlaid spots showing expression profiles characterised by the obtained clusters.

Users can interactively select genes of interest, adjust visualization parameters such as alpha transparency and point size, and choose between different colour schemes to enhance the representation of gene expression levels. The atlas provides an intuitive platform for investigating spatial gene expression patterns and is publicly available online at https://adrenal-spatall-transcriptomic.shinyapps.io/Adrenal/ as of November 11, 2024.

### Gene ontology and pathway enrichment analysis of marker genes

To elucidate the biological functions and pathways associated with the differentially expressed genes (DEGs) identified in each adrenal gland cluster, we performed Gene Ontology (GO) and Kyoto Encyclopedia of Genes and Genomes (KEGG) pathway enrichment analyses. DEGs with an adjusted *p*-value less than 0.05 and an average log2 fold change greater than 1 were considered significant and included in the analysis. The top ten enriched GO terms for each cluster were identified and visualized using bubble plots (**Figure 2**), where the size of each bubble represents the number of genes associated with a term, and the colour intensity reflects the statistical significance. Using the GO BP DIRECT reference database, the most significantly enriched pathway in the ZG cluster was the Wnt signaling pathway (adjusted *p*-value = 3.86×10^−8^), involving genes such as *Apcdd1* (adenomatosis polyposis coli down-regulated 1), *Ccnd1* (cyclin D1), *Fzd6* (frizzled class receptor 6), *Lef1* (lymphoid enhancer binding factor 1), and *Wnt4* (Wnt family member 4). Other highly enriched GO terms included negative regulation of canonical Wnt signaling pathway and canonical Wnt signaling pathway, indicating a pivotal role of Wnt signaling in ZG function. Additionally, processes related to animal organ morphogenesis and cell adhesion were enriched. For the ZF cluster, the top enriched GO term was the lipid metabolic process (adjusted *p*-value = 1.34×10^−3^), with genes such as *Abhd3* (abhydrolase domain containing 3), *Acsl4* (acyl-CoA synthetase long chain family member 4), *Cyp11a1* (cytochrome P450 family 11 subfamily A member 1), and *Lpin3* (lipin 3) playing crucial roles. The fatty acid metabolic process was also significantly enriched, aligning with the ZF’s role in glucocorticoid synthesis and lipid metabolism. The steroid biosynthetic process term was notably enriched (adjusted *p*-value = 1.76×10^−1^), involving key steroidogenic genes such as *Cyp11a1, Cyp11b1* (cytochrome P450 family 11 subfamily B member 1), *Cyp21a1* (cytochrome P450 family 21 subfamily A member 1), and *Star* (steroidogenic acute regulatory protein). In the ZX cluster, the steroid biosynthetic process was the most significantly enriched term. Lipid metabolic process and cholesterol metabolic process were also highly enriched. The medulla cluster, originating from the neuroectodermal lineage of the adrenal medulla, showed significant enrichment in neural-dependent processes. The top GO term was nervous system development (adjusted *p*-value = 3.98×10^−20^), involving genes such as *Ascl1* (achaete-scute family bHLH transcription factor 1), *Gata3* (GATA binding protein 3), *Ntrk1* (neurotrophic receptor tyrosine kinase 1), and *Slit2* (slit guidance ligand 2). Other enriched terms included modulation of synaptic transmission, ion transmembrane transport, and axon guidance, reflecting the medulla’s role in catecholamine synthesis and neuronal function. The complete set of enriched terms, referencing all analyzed GO categories and KEGG pathways, is provided in Supplementary File 2.

**Figure 2.**
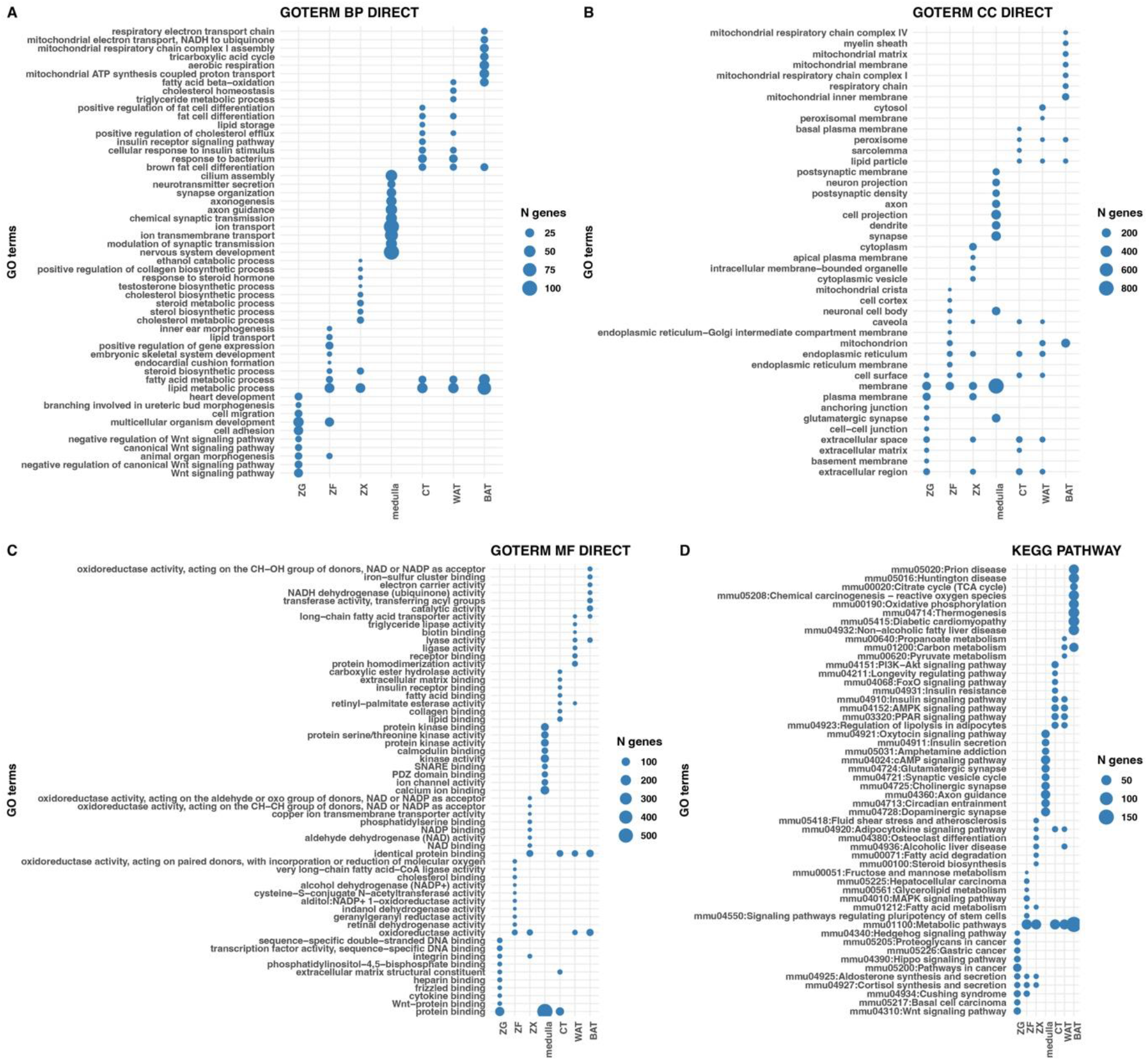
Bubble plots of the top enriched GO terms and KEGG pathways for each cluster. X-axis represents the different clusters corresponding to adrenal gland regions or cell types (ZG, ZF, ZX, medulla, CT, WAT, BAT). Y-axis corresponds to the enriched GO terms and KEGG pathways, grouped by ontology categories: A) Biological Processes (GO BP), B) Molecular Functions (GO MF), C) Cellular Components (GO CC) and D) KEGG pathways. Bubble Size indicates the number of DEGs associated with each term.

### Spatial deconvolution and characterization of adrenal gland spots heterogeneity

Next, we utilized the Cell-type Assignment by Regression on Distances (CARD) algorithm to achieve spatial deconvolution of complex mixtures of cell types, estimating the relative proportions of each cell type across the adrenal gland tissue (**Figure 3**). This analysis employed a reference-free approach and integrated single-cell transcriptomics data to generate a comprehensive list of biomarkers for specific adrenal cell types and auxiliary components, such as fibroblasts, endothelial cells, and white and red blood cells (Supplementary File 3). We further analysed cell-type composition by performing spatial correlation modelling to refine tissue maps, enhancing the resolution of cell-type distributions at specific spatial locations. This approach validated and extended the cluster-based classification identified earlier in the study. Notably, we successfully characterized and distinguished adipose tissue types in the adrenal glands, specifically BAT and WAT. Additionally, our analysis delineated distinct zones within the adrenal cortex, including the ZG, ZF, and ZX. A strong correlation was observed between the medulla and red blood cells (RBCs), consistent with the anatomical structure of the adrenal gland and the vascularized nature of the medulla. Key biomarkers with the highest expression levels for each cluster were identified (**Figure 3C**). In the ZG, markers such as *Dab2* (disabled 2, mitogen-responsive phosphoprotein) and *Cyp11b2* (cytochrome P450 family 11 subfamily b polypeptide 2) were highly localized, while *Prss35* (serine protease 35) and *Akr1b7* (aldo-keto reductase family 1 member B7) were prominently expressed in the ZF region. The ZX was marked by the presence of *Abcb1b* (ATP-binding cassette sub-family B member 1B), *Ly6m, Mapk13* (mitogen-activated protein kinase 13), and *Sbsn*. In the medulla, *Npy* (neuropeptide Y), *Th* (tyrosine hydroxylase), and *Rab3c* (RAB3C, member RAS oncogene family) were identified as key genes associated with catecholamine synthesis. These findings underscore the unique molecular architecture of each adrenal gland component. The spatial localization of distinct markers provided further insights into the physiological functions of the adrenal zones and their specialized roles in steroid biosynthesis and neural activity. The CARD analysis not only reinforced the spatial and molecular delineation of adrenal gland components but also illuminated the complexity of various cell types across the gland, offering a robust framework for understanding adrenal gland biology.

**Figure 3.**
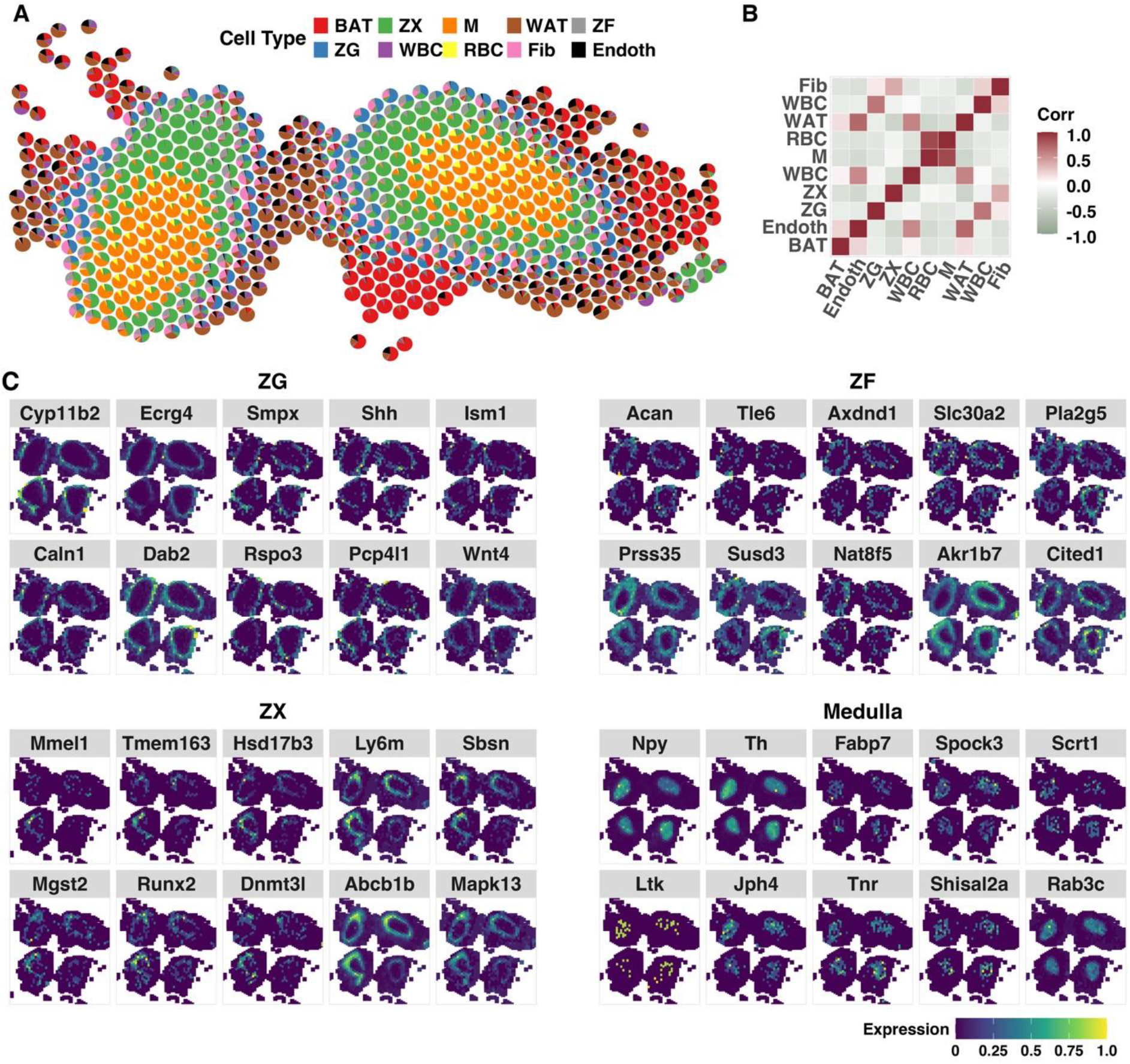
Spatial distribution of identified biomarkers for adrenal gland components. The Cell-type Assignment by Regression on Distances (CARD) analysis based on the list of markers from single-cell data within mouse adrenals. A) The cell proportion matrix mapped onto the adrenal gland spatial transcriptomics analysis. Each cell type is colour-coded, revealing distinct clustering patterns. B) Cell type proportion correlation of selected groups of cells. C) Visualizations of the marker gene expression in ZG, ZF, ZX and Medulla with the spatial locations within the tissue. Fib - fibroblasts, Endoth - endothelial cells, WBC - white blood cells, RBC - red blood cells.

### Pseudotime trajectory analysis confirmed the centripetal differentiation within the adrenal cortex

Using spatial transcriptomic expression data, we investigated the centripetal differentiation within the mouse adrenal cortex through pseudotime analysis using the Monocle 3 library (Figure 4). For this analysis, only spots corresponding to the CT capsule and adrenal cortex zones were extracted and subjected to trajectory and diffusion pseudotime analysis to infer the developmental progression of cortical cell populations. Our trajectory analysis confirmed the centripetal differentiation within the adrenal cortex, revealing a clear progression of cells transitioning through adrenal cortex zones over pseudotime (Figure 4B). Specifically, the differentiation and transition sequence of adrenal cell types begins at the CT capsule (earliest pseudotime), progresses to the ZG, then to the ZF, and finally to the ZX, which represents the most mature stage of differentiation (Figure 4B, C). To explore functional differences between these zones, we analysed the expression of key genes (Figure 4D). The sequence of cell transitions aligns with the known steroidogenic activities of zone-specific markers. For instance, genes such as *Cyp11b2* and *Ecrg4*, which are predominantly expressed in the ZG, exhibit expression patterns consistent with early pseudotime stages. Conversely, genes associated with steroidogenesis in deeper cortical layers, such as *Cyp11a1* (cytochrome P450 family 11 subfamily A member 1), *Srd5a2* (steroid 5-alpha-reductase 2), and *Cyp11b1* (cytochrome P450 family 11 subfamily B member 1), demonstrate higher expression in more mature zones, supporting the proposed differentiation sequence.

**Figure 4.**
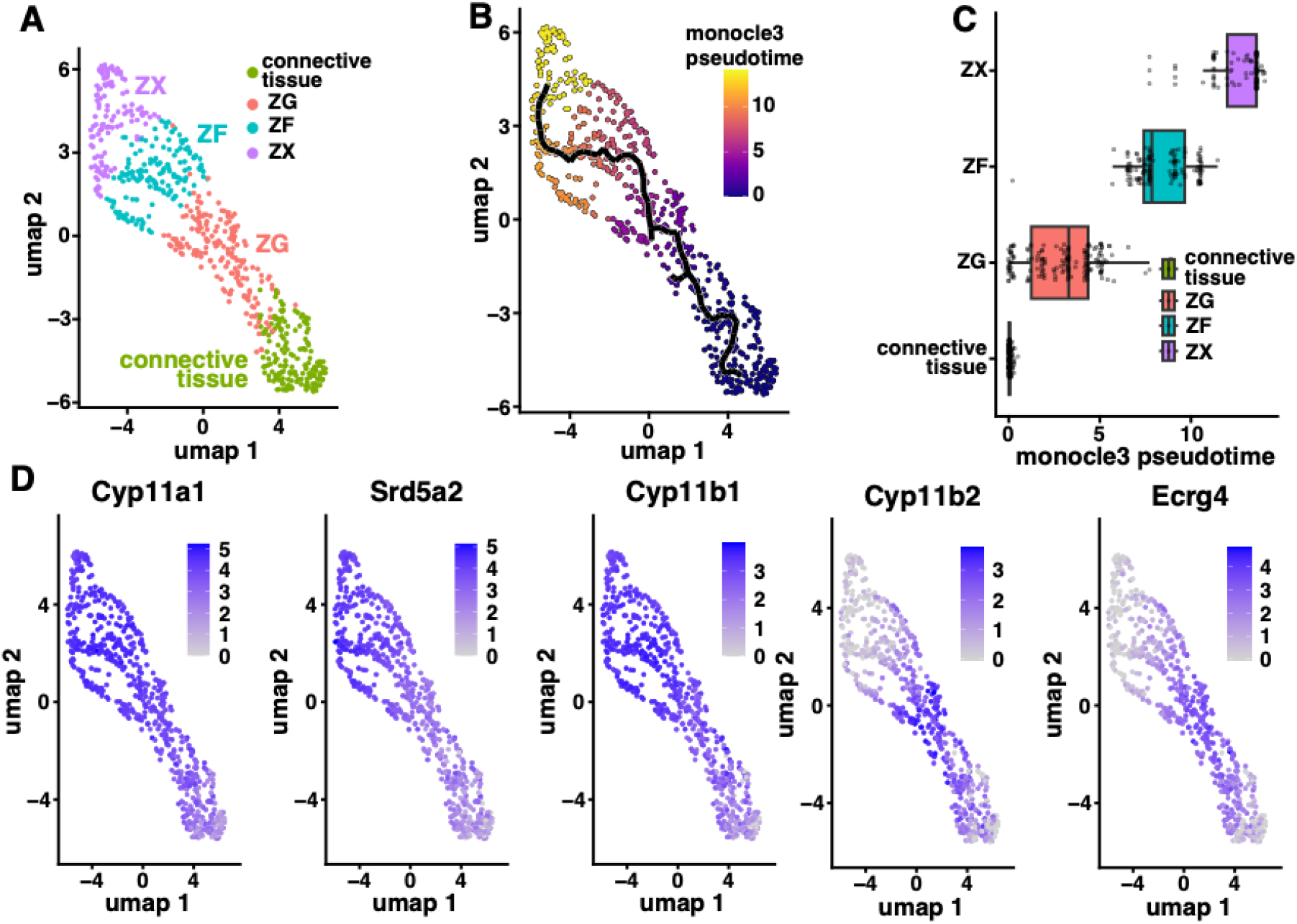
Pseudotime (trajectory) analysis of spots within the adrenal cortex and connective tissue capsule. A) UMAP dimensionality reduction with visualisation of the spots in the adrenal cortex and connective tissue clusters. B) Trajectory graph generated from the UMAP dimensionality reduction of the spots coloured by pseudotime, according to the colour scale representing the start and end states of the trajectory. The colour gradient from purple to yellow indicates pseudotime progression. C) The range of pseudotime in the analysed clusters (connective tissue, ZG, ZF, ZX) ordered by mean pseudotime value. D) Examples of genes whose expression changes along pseudotime in analysed spots.

### Cell-cell communication network in the mouse adrenal gland

To investigate intercellular communication within the mouse adrenal gland, we performed CellChat analysis on spatial transcriptomics data. This approach allowed us to predict ligand-receptor interactions between different regions of the adrenal gland, including the ZG, ZF, an undefined cortical ZX, medulla, CT, WAT, and BAT. Our analysis revealed several key signalling pathways mediating communication predominantly within the adrenal cortex (Figure 5), where cumulative probabilities between groups are presented. For single representative ligand-receptor pairs, we applied a probability threshold of 0.005 to focus on the most significant interactions. Canonical WNT signalling involving Wnt4 and its receptor complex *Fzd3-Lrp6* was prominent within and between cortical zones. Significant interactions were observed between ZG and ZF, ZG and ZX, as well as within ZG and ZF themselves (probability range: 0.0083–0.0173). Non-canonical WNT (ncWNT) signalling via *Wnt5a-Fzd3* was also significant within the cortex. Interactions were detected between ZG and ZF, ZG and ZX, ZF and ZX, and within ZG, ZF, and ZX (probability range: 0.0062–0.0186). The Hedgehog signalling pathway emerged as a significant mediator of communication within the cortex and from cortex to connective tissue. Sonic Hedgehog (Shh) produced by ZG and ZF interacts with its receptor *Ptch1* expressed in ZF, ZX, and CT (probability range: 0.0069–0.0281). Insulin-like growth factor 2 (*Igf2*) expressed in the medulla interacts with its receptor *Igf2r* in ZF and ZX (probability range: 0.0067–0.0139), suggesting that medullary *Igf2* may influence on cortical cell proliferation, differentiation, or function via paracrine mechanisms. Macrophage migration inhibitory factor (*Mif*) interacted with *Ackr3* across cortical zones and between the cortex and medulla (probability range: 0.0119–0.1174). Galectin-9 (*Lgals9*) interactions with *P4hb* were significant within cortical zones and between the cortex and medulla (probability range: 0.0170–0.0644). Galectin signalling is associated with cell adhesion, apoptosis, and immune regulation, indicating that it may contribute to cortical cell maintenance. Endothelin signalling via *Edn3-Ednrb* from the medulla to cortical zones, particularly to ZX (probability: 0.0129), suggests a role in vasoconstriction and modulation of cortical blood flow. Our analysis identified significant interactions involving galanin (*Gal*) and its receptor *GalR1*, predominantly within the medulla itself (probability: 0.0091), suggesting a role in medullary function. Significant interactions involving *Npy* and its receptor *Npy2r* were observed within the medulla (probability: 0.1449), indicating a strong autocrine signalling role of *Npy* in the medulla.

**Figure 5.**
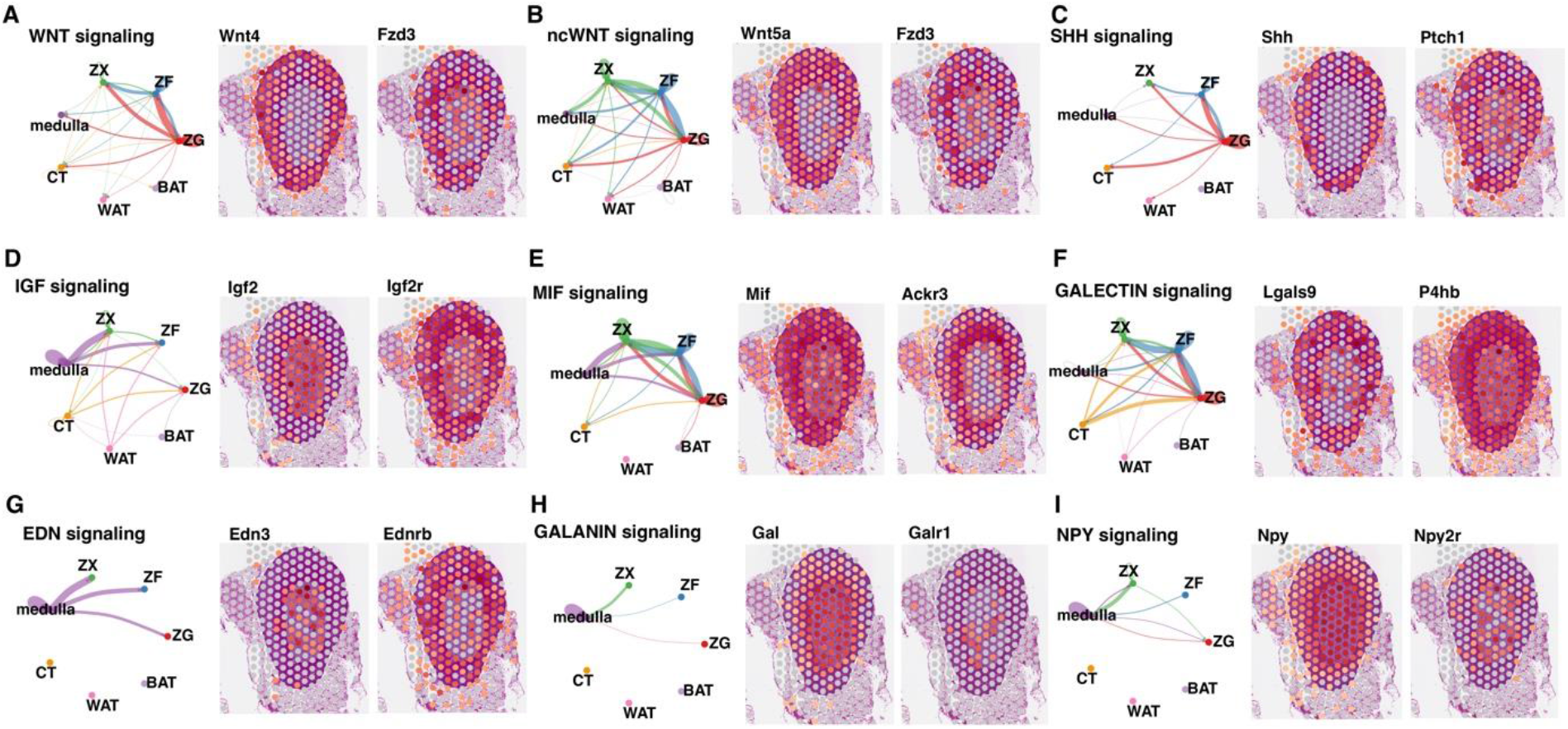
Inferred communication networks of individual L-R (ligand-receptor) pairs associated with signalling at A) Wnt, B) non canonical WNT, C) SHH, D) IGF, E) MIF, F) GALECTIN, G) EDN, H) GALANIN and I) NPY pathways. The left panel shows circle plots illustrating communication strength between or within the analyzed groups, where the thickness of the connecting lines represents the cumulative communication probability. The right panel displays the spatial expression of ligand (left side) and receptor (right side) pairs with the highest communication probability between analysed groups for each specific signalling pathway.

## Discussion

To date, research on the adrenal gland has focused on gene expression profiling, hormone and metabolite measurements, immunohistochemical analyses, genetic studies, and cell culture models. While these approaches have been insightful, they have not yet captured the complete molecular landscape of dynamic cellular changes within the adrenal gland. Histological and molecular biology methods such as laser microdissection or omics studies, did not allow for complete analysis of individual adrenal cellular components or their surrounding structures. As a result, previous studies attempting to address questions related to adrenal function have been hampered by errors due to difficulties in obtaining pure adrenal cellular components. Often, despite efforts to maintain precision, isolated structures suffered from errors and contamination, which could affect the purity of the target population. Therefore, in the absence of unique molecular markers, studies of adrenal structure and function can be a serious scientific challenge.

In this study, we utilized spatial transcriptomics to elucidate the molecular landscape of the CD1 IGS adult mouse adrenal gland, identifying genetic markers associated with adrenal zonal identification and providing broader insights into the dynamic cellular interactions within this complex organ. Furthermore, our research provides novel insights into the molecular basis of adrenal histological heterogeneity. We achieved this by examining the spatial distribution of gene expression signatures and their association with distinct histological compartments. Our analysis identified five clusters that align with adrenal gland histology (ZG, ZF, ZX, medulla, and connective tissue capsule) and two distinct clusters of adipose tissue (brown and white).

In our studies, we have shown that each of the defined zones is characterised by a unique molecular signature that is directly related to the physiological functions of the adrenal gland. In the case of the ZG, we identified genes such as *Cyp11b2, Ecrg4*, closely linked to aldosterone synthesis, or *Rspo3* (R-spondin 3), essential for the renewal of the adrenal cell pool in the ZG, whereas genes related to glucocorticoid production, such as *Acan* (Aggrecan) or *Susd3*, are precisely present in the ZF (Aiba & Fujibayashi, 2011; Neville, 1982; Porzionato *et al*, 2015; Vidal *et al*, 2016). Some of the genes mentioned are known from the literature(Aiba & Fujibayashi, 2011; Neville, 1982; Porzionato *et al*., 2015; Vidal *et al*., 2016), confirming the results of our analyses, which provided a comprehensive resolution by describing the specific cell regions in the adrenal gland. However, in the case of genes such as *Frmpd4* (FERM And PDZ Domain Containing 4), *Oca2* (P gene), *Sphkap* (SPHK1 Interactor, AKAP Domain Containing) for the ZG or *Cited1, Nat8f5* (N-acetyltransferase 8) for the ZF, although the presence of some of this gene has been confirmed in adrenal tissue (Splittstosser *et al*, 2019; Val *et al*, 2007; Xu *et al*, 2016), they have not been proposed as prominent markers useful in distinguishing individual zones of the adrenal cortex. This discovery may point to a new role for these genes in the functioning of individual zones of the adrenal cortex and may open a new avenue of research to understand the influence of these markers on the mechanisms of adrenal physiology. However, this finding needs to be confirmed in a larger cohort. The ZX zone was characterised by increased expression of *Ly6m, Sbsn* or *Abcb1b*, which are closely related to androgen production (El Wakil *et al*, 2013; Gannon *et al*., 2019; Hershkovitz *et al*, 2007; Huang & Kang, 2019; Lopez *et al*, 2021). Interestingly, the presence of the ZX region in sexually mature male mice and suggest that the lack of ZX involution may be a characteristic feature of this species. Overexpression of potential ZX genes has not yet been characterised in this region but may be necessary in adaptive processes related to changes in the metabolic or protective requirements of the ZX zone prior to its involution. Both strain-specific differences and ZX development may have implications for understanding adrenal development and dystrophy. Next, at the medullary region we indicate the enrichment in genes like *Npy* or *Th*, as well as *Jph4, Gnb3, Mast1*, and *Tnr*. All genes are associated with neuroendocrine function and primary source of catecholamines, reflecting the medulla’s role as a critical for the body’s acute stress response. While *Npy* and *Th* are typically expressed in the adrenal medulla, the remaining proposed genes have not yet been associated with the adrenal medulla. One of these is the *Jph4* gene, which, like *Mast1*, is mainly expressed in neurons and cells of the nervous system(Hogea *et al*, 2021; Jing *et al*, 2020). This discovery furthers our understanding of chromaffin cells and their potential interactions with the surrounding adrenal cortex. Furthermore, two clusters were identified that are abundant in the surrounding area of the adrenal tissue. These clusters are characterised by the presence of *Ucp1* and *Cidec*, respectively, and are known as BAT and WAT (Fujita *et al*, 2021; Shen *et al*, 2016). The potential interactions between these adipose tissues and the adrenal cortex highlights a potential regulatory relationship, possibly involving steroid hormone production and lipid metabolic regulation. BAT is known for its role in thermogenic capacity and might exert an influence on adrenal function under conditions of cold-induced stress (Jiang *et al*, 2024; Witt *et al*, 2023). Meanwhile, WAT is involved in energy storage and the secretion of adipokines, which may potentially affect adrenal steroidogenesis. An understanding of these relationships is crucial for comprehending the broader implications of adrenal function in the context of energy homeostasis and stress responses.

The results of pseudotime trajectory analysis corroborate the centripetal differentiation model of the adrenal cortex, demonstrating a progression of cellular differentiation from the outer connective tissue capsule through the ZG and ZF to the ZX. In this model, cells undergo dynamic migration towards the inner cortex, accompanied by shifts in molecular and functional properties. Our findings corroborate those of previous lineage-tracing studies, which have suggested the presence of progenitor cells between the adrenal capsule and ZG (Altieri *et al*, 2024; Freedman *et al*., 2013; Pihlajoki *et al*., 2015; Vinson, 2016). These progenitor cells give rise to differentiated adrenocortical cells that migrate inward. Furthermore, our findings have been supported by analysing a pseudotime-dependent expression of key markers, including *Cyp11a1, Srd5a2, Cyp11b1, Cyp11b2*, and *Ecrg4*, which mark different stages of steroidogenic differentiation within the adrenal cortex. More recently, single-cell RNA sequencing (scRNA-seq) and spatial transcriptomics have provided high-resolution insights into adrenal cell differentiation in both mouse and human adrenals (Del Valle *et al*, 2023; Fu *et al*, 2023; Iwahashi *et al*, 2024). In this context, our research confirms the centripetal differentiation model within the adrenal cortex, highlighting the gradual transition of cell populations from the connective tissue capsule through distinct cortical zones. Furthermore, the expression dynamics of steroidogenic genes underscore the functional specialization of each zone, reinforcing the role of spatial and pseudotime analyses in understanding adrenal gland development and function.

Using CellChat ligand-receptor interaction network in spatial format, we demonstrated that expression of specific genes is arranged in characteristic patterns corresponding to several adrenals signalling pathways in each of adrenal components. Developmental signalling pathways regulate the process of cellular pluripotency, differentiation, and patterning in various tissues(Lerario *et al*., 2017; Pihlajoki *et al*., 2015; Tai & Shang, 2023). In the adult organism, some of these pathways play a pivotal role during the rapid growth phase of adrenal development. By elucidating and meticulously mapping a complex network of ligand-receptor interactions within the mouse adrenal gland, we highlight an insight into the functional specialisation of each adrenal zone. We identified several key pathways, including WNT, SHH, IGF, and GALECTIN. These pathways facilitate both inter-zone and intra-zone communication, thereby highlighting potential mechanisms for the regulation, maintenance and response of the adrenal gland to physiological demands. For example, the activation of WNT and SHH signalling pathways confirm a crucial role for these pathways in coordinating adrenal development and zonal differentiation(Basham *et al*, 2019; Walczak & Hammer, 2015; Walczak *et al*, 2014). Moreover, the disruption of WNT signalling has also been associated with adrenocortical carcinoma development(Borges *et al*, 2020; Tai & Shang, 2023). Understanding the molecular mechanism underlying adrenal zonation and differentiation may lead to expand strategies for treating adrenal diseases. Meanwhile, the persistence of the X-zone provides the opportunity to explore its functional significance in adult mice, including potential roles in androgen synthesis and adrenal remodelling. The GALECTIN communication by *Lgals9* might modulate immune response and cellular adhesion(Golden-Mason *et al*, 2013). Above suggest the potential of ZX in adrenal immune interactions or stress response mechanisms. Similarly, in the adrenal medulla we identified the NPY signalling which play a pivotal role in regulating neuroendocrine functions and stress responses, thereby underscoring the distinct regulatory requirements of the adrenal medulla (Descamps *et al*, 2001; Forander *et al*, 2000; Goedert *et al*, 1983). Identifying ligand-receptor interactions between distinct adrenal zones suggests the existence of a complex regulatory network that highlights the adrenal gland’s role as a central hub for integrating endocrine and neuroendocrine signals in response to stress.

It is acknowledged that the findings presented herein have limitations. Despite the advanced spatial resolution of the Visium platform, single-cell heterogeneity may still need to be fully captured. Furthermore, the analysis was conducted using a single developmental stage of the mouse. Therefore, future studies should include multiple ages and sexes to enhance the generalizability of the observed findings. A closer examination of the adrenal development process and investigation of the potential sexual dimorphism and age-related differences in adrenal renewal processes are essential for a comprehensive understanding of the subject matter.

It should be emphasised that, to our knowledge, this is the first study using Visium Cyt.Assist technology carried out on the adrenal glands of animals in which the ‘transient structure (ZX)’ occurs. Our results provide, for the first time, a fundamental map of the spatial transcriptomic organisation of the adult mouse adrenal gland, providing a further understanding of adrenal function and regulation. By identifying novel genetic markers associated with adrenal zonal differentiation and confirming the centripetal differentiation model, we contribute to a deeper understanding of adrenal biology. The detailed spatial map may facilitate the identification of new therapeutic targets for adrenal-related disorders, as well as provide insights into the cellular interactions underlying the adrenal gland’s response to metabolic and environmental demands. Given the similarities in adrenal cortex structure and function between mice and humans, our findings using the mouse model may have significant implications for understanding human adrenal disorders and developing potential therapeutic interventions.

## Material & Methods

### Ethical statement

Animal experiments involving mice (Mus musculus) were conducted in compliance with Directive 2010/63/EU on the protection of animals used for scientific purposes and the Polish Act on the Protection of Animals Used for Scientific or Educational Purposes. The study was approved by the Local Ethics Committee for Animal Studies in Poznan, Poland (protocol No. 27/2023, dated April 17, 2023). Animal welfare was regularly monitored by evaluating their activity levels, behaviour, water and food consumption, as well as the condition of their fur and stool.

### Mice

Sexually mature male CD-1® IGS mice, 10 weeks old with a body weight of approximately 37 g were used in this study. The experiment was approved by the Local Ethical Commission in Poznan (Decision no. 27/2023, dated April 28^th^, 2023). All mice were bred by the Centre for Advanced Technology (Adam Mickiewicz University, Poznan). Mice were housed in colony cages in a pathogen-free environment with temperature maintained at 21-23°C and relative humidity between 55-60% and under a 12h light/12 h dark cycle. All mice were given standard chow and water *ad libitum*. The mice were euthanized by decapitation and the harvested adrenals were immediately preserved in 10% buffered formalin for further analysis.

### Tissue processing and Spatial transcriptomics

Regions of interest (ROIs) of the mouse adrenal glands were selected based on morphology (expert review by M.B.), orientation (based on H&E staining) and quality assessment of the extracted RNA by DV200 that was obtained using an Agilent 2100 Bioanalyzer. Next, the FFPE blocks were sectioned at 5 μm using a microtome (Leica RM2255). The isolation of total RNA from FFPE tissue blocks was done by using the RNeasy FFPE kit (73504, Qiagen).

FFPE Visium Spatial Gene Expression (10x Genomics) was performed on CytAssist instrument, following the manufacturer’s protocol. Briefly, the tissue sections were placed on SuperFrost™ Plus Microscope Slides (#22-037-246, FisherScientific) and dried overnight. Then, the slides underwent incubation at 60°C for a 2 hours, followed by deparaffinization. Subsequently, the sections were stained with H&E and imaged at 20× magnification in brightfield using Grundium Ocus 20 (Grundium). For the H&E-stained sections, decrosslinking was carried out immediately and then probe panels covering the entire mouse transcriptome were incorporated. After the probe pairs were hybridized with their target transcripts, the slides were placed on a Visium CytAssist instrument for permeabilization and RNase treatment. After transcript capture, the Visium Library Preparation protocol from 10x Genomics was performed. Four cDNA libraries were diluted and pooled to a final concentration of 650 pM (20 μl volume) and sequenced on P1 flow cell of an Illumina NextSeq 2000.

### Data processing

All datasets have been deposited in the Gene Expression Omnibus (GEO) database under accession number GSE283302. Demultiplexing of the sequencing reads was performed with Illumina bcl2fastq (v2.20). Visium Spatial Gene Expression analysis was performed on a local server (Ubuntu 22.04.1 LTS) with Space Ranger software (version 2.0.1) by 10x Genomics. Sequencing data for each tissue sample (in FASTQ format) and corresponding capture area image (TIFF, QPTIFF, or JPEG format) were analysed by running the Space Ranger count pipeline with slide details and reference transcriptome, for mouse, as arguments. In the case of analysis for FFPE samples, the count pipeline required a probe set reference CSV file. Appropriate probe set files and available mouse (mm10) reference datasets required for Space Ranger were downloaded from 10x Genomics. For multi-sample analysis, results from each run were aggregated into single output files by spaceranger aggr pipeline with aggregation CSV file as argument.

### General data analysis

The raw data from Space Ranger were processed and analysed using R (version 4.1.2; R Core Team 2021) and the Seurat package for the pre-processing workflow, which included normalisation, scaling, clustering, and dimensionality reduction (Hao *et al*, 2021; Hao *et al*, 2024). The “Load10X_Spatial” function was employed to load the spatial gene expression data along with the associated high-resolution tissue image. To account for differences in image resolution, scale factors were adjusted to match the image’s dots per inch (DPI) to 300, ensuring accurate spatial mapping. Gene expression normalization was performed using the “NormalizeData” function with the “LogNormalize” method and a scaling factor of 10,000. Highly variable features were identified using the “FindVariableFeatures” function with the “vst” selection method, retaining the top 2,000 variable genes. All genes were subsequently scaled using the “ScaleDatafunction” to standardize the expression levels. General expression profile of individual genes was also displayed as spots with expression profiles on a histological slide from Loupe Browser 7 software (10X Genomics).

### Dimensionality reduction and clustering

Principal component analysis (PCA) was conducted using the RunPCA function on the previously identified variable features. The number of components for the analysis was determined by evaluating dimensionality through the JackStraw resampling method provided in the ‘Seurat’ package (Hao *et al*., 2021; Hao *et al*., 2024). Spot clustering was performed using the K-nearest neighbors (KNN) algorithm combined with Jaccard similarity, as implemented in the ‘FindNeighbors’ function. Subsequently, the data were input into the ‘FindClusters’ function, which established clusters based on a resolution parameter that controls the level of clustering detail. After generating clusters at multiple resolutions, the optimal clustering resolution was selected based on the stability and biological relevance of the identified clusters. Uniform Manifold Approximation and Projection (UMAP) was utilized for dimensionality reduction and visualization using the RunUMAP function on the first 10 principal components. Clusters were visualized in two-dimensional space to assess the separation and distribution of cell populations. Data visualization was performed using the ggplot2, patchwork, and cowplot packages (Pedersen, 2024; Wickham, 2016; Wilke, 2024).

### Cluster annotation and marker gene identification

Clusters were annotated based on the expression of known marker genes and prior biological knowledge. Specifically, clusters were labeled to represent distinct adrenal gland regions and tissue types, including ZG, ZF, ZX, medulla, WAT, and BAT. Cluster identities were updated accordingly using the Idents function in Seurat. Differentially expressed genes (DEGs) for each cluster were identified using the “FindAllMarkers” function. The analysis selected only genes that were positively expressed with an average log2 fold change threshold of 1, detected in at least 25% of cells within a cluster and adjusted p-values less than 0.05. The top 10 marker genes for each cluster were visualized as heatmap using “DoHeatmap” function. The list of potential markers was also saved to an xlsx file using the openxlsx package (Schauberger & Walker, 2024).

### Gene Ontology and Pathway Enrichment Analysis

Functional enrichment analysis was performed using the RDAVIDWebService package to interface with the Database for Annotation, Visualization, and Integrated Discovery (DAVID) (Dennis *et al*, 2003). Mouse gene symbols were converted to Entrez Gene IDs using the biomaRt package to obtain require annotation for ontology analysis(Durinck *et al*, 2009). Enrichment analyses for Gene Ontology (GO) terms—including biological processes (BP), molecular functions (MF), and cellular components (CC)—as well as KEGG pathways were conducted. Terms with an adjusted *p*-value less than 0.05 were deemed significant. Ten ontology groups with the lowest adj *p* value for each of clusters were visualized as bubble plots generated by the ggplot2 package (Wickham, 2016).

### Cell-cell communication analysis

The CellChat package was used to analyse and visualise the cell-cell communication network for a spatial transcriptomics dataset, following the provided manual(Jin *et al*, 2024). For the previously created Seurat object, spatial transcriptomic data and spatial coordinates were retrieved using the ‘GetTissueCoordinates’ function. The spatial factors were computed through pixel-to-micrometer conversion by determining the pixel ratio of the theoretical spot size (65 um) to the number of pixels covering the diameter of the spot size in the full-resolution image provided in the ‘scalefactors_json.json’ file. The obtained spatial coordinates and spatial factors were used as arguments to the ‘createCellChat’ function to construct the CellChat object. The analysis utilized the CellChatDB.mouse database, focusing on secreted signaling pathways. Overexpressed genes and interactions were identified using the identifyOverExpressedGenes and identifyOverExpressedInteractions functions, respectively. Communication probabilities were computed using the computeCommunProb function with the truncated mean method, considering spatial distances with an interaction range of 250 μm and a scaling factor of 0.01. Contact-dependent signalling within a range of 100 μm was also included. The cell-cell communication network was aggregated by either counting the number of links or summarising the communication probability. Inferred communication networks of individual L-R (ligand-receptor) pairs associated with that signaling pathway were visualized as circle plot. The expression profile of ligand-receptor pairs with the strongest effect on communication probability was shown on histological slides.

### Pseudotime (trajectory) analysis

To explore the developmental trajectories and lineage relationships among adrenal gland cell populations, pseudotime analysis was performed using the Monocle 3 package, following the guidelines provided in the Monocle 3 manual (Cao *et al*, 2019; Trapnell *et al*, 2014). Specifically, only spots corresponding to CT, ZG, ZF, and ZX were extracted from the entire dataset for this analysis. Preprocessing was conducted in the same manner as for UMAP dimensionality reduction to ensure consistency. This included normalization using the NormalizeData function with default parameters, identification of highly variable features using FindVariableFeatures, data scaling via ScaleData, and principal component analysis (PCA) using RunPCA with the top 10 principal components. Clustering information and UMAP embeddings from Seurat were transferred to the Monocle 3 object. To determine the optimal trajectory, the learn_graph function in Monocle 3 was utilized without partitioning (use_partition = FALSE). This function constructs a principal graph that captures the global structure of the data and models the developmental pathways. The analyzed spots were then sorted based on their progression through the developmental program using the order_cells function, with connective tissue cells designated as the root cells. This rooting assumes that the developmental trajectory originates from the connective tissue, allowing for the assessment of differentiation progression into other zones (ZG, ZF, ZX). Pseudotime values were computed for each cell, representing their position along the developmental trajectory. The obtained results were presented as UMAP graphs and box plots. UMAP plots displaying cells coloured by pseudotime and cluster identities were generated using the plot_cells function in Monocle 3. Box plots illustrating the distribution of pseudotime values across the different clusters (CT, ZG, ZF, ZX) were created using the ggplot2 package

### Cell-type Assignment by Regression on Distances (CARD) -deconvolution analysis

To deconvolute spatial gene expression profiles and estimate cell-type proportions within the mouse adrenal gland tissue sections, we employed the Cell-type Assignment by Regression on Distances (CARD) package(Ma & Zhou, 2022). To prepare the data for CARD analysis, the Seurat object was converted into a SpatialExperiment object using the SpatialExperiment package. A reference-free approach was adopted for the deconvolution analysis, utilizing a list of marker genes generated from scRNA-seq data downloaded from the Gene Expression Omnibus (GEO) database (accession number: GSE162238(Hanemaaijer *et al*, 2021)) and from markers described in literature. The marker genes were identified based on their specificity to particular cell types within the adrenal gland and were curated into a structured list. Deconvolution was performed using the CARD_refFree function, which estimates cell-type proportions in each spatial spot based on the marker gene list and spatial gene expression data. The analysis inferred the distribution of various cell types, including ZG, ZF, ZX, medulla, fibroblasts, endothelial cells, white blood cells, red blood cells, and adipose tissue types. The results from the CARD analysis were visualized using several functions within the CARD package and additional R packages. Spatial pie charts representing the estimated cell-type proportions across the tissue section were generated using the CARD.visualize.pie function, with customized color palettes from the RColorBrewer package. To examine the expression patterns of top marker genes for each cell type, the CARD.visualize.gene function was used. This function created high resolution spatial feature plots for selected marker genes, allowing visualization of their spatial distribution within the tissue. The top 10 marker genes for each cell type were selected based on their expression levels and specificity. In current paper only markers for adrenal ZG, ZF, ZX and medulla are presented. Correlation analysis between inferred cell-type proportions and cluster identities from the Seurat analysis was performed by integrating the CARD results with the cluster metadata. Mean cell-type proportions were calculated for each cluster using the dplyr package and visualized by ggplot2 package (Wickham, 2016).

### Interactive Visualization of Spatial Transcriptomics Data

To facilitate the exploration and interactive visualization of the spatial transcriptomics atlas of the mouse adrenal gland, a Shiny web application was developed. The application was based on protocols for scRNA-seq and spatial gene expression integration and interactive visualization. The Shiny application was developed using the Shiny package(Chang *et al*, 2024), with additional packages including ggplot2, Seurat, RColorBrewer(Neuwirth, 2022), miniUI (Cheng, 2018), and grid(Team, 2024). Color palettes were precomputed using the colorRampPalette function from the RColorBrewer package to enhance visualization flexibility. The user interface (UI) was constructed using the miniPage and miniContentPanel functions. The application allowed users to interactively select any gene present in the dataset to visualize its spatial expression pattern across the adrenal gland tissue section. Adjustments to alpha transparency and point size provided enhanced visualization of overlapping data points and control over plot aesthetics.

---

**Supplementary File 1:** Marker genes identified in the spatial transcriptomic analysis of the mouse adrenal gland for zones ZG, ZF, ZX, medulla, CT, WAT, and BAT. The dataset includes the following columns: p_val (raw p-value), avg_log2FC (average log_2_ fold change in expression), pct.1 (percentage of spots within the cluster where the gene is detected), pct.2 (highest percentage of spots in other clusters where the gene is detected), p_val_adj (adjusted p-value), cluster (cluster name), and gene (gene symbol).

**Supplementary File 2:** Supplementary File 2: Results of Gene Ontology (GO) and KEGG pathway enrichment analyses for the specified clusters. The file contains four sheets: GO BP (enriched GO Biological Process terms), GO MF (enriched GO Molecular Function terms), GO CC (enriched GO Cellular Component terms), and KEGG (enriched KEGG pathways). Each sheet includes the following columns: Category (the GO term category or KEGG pathway), Term (the specific GO term or KEGG pathway name), Count (number of genes associated with the term), PValue (raw p-value of the enrichment analysis), Genes (list of gene symbols associated with the term), List.Total (total number of genes in the input list), Pop.Hits (number of genes associated with the term in the background population), Pop.Total (total number of genes in the background population), Fold.Enrichment (ratio of observed gene count to expected count, i.e., enrichment score), Bonferroni (Bonferroni-corrected p-value), Benjamini (Benjamini-Hochberg adjusted p-value), FDR (false discovery rate), and Cluster (name of the cluster).

**Supplementary File 3:** Marker genes identified for deconvolution across various clusters. The file contains gene symbols associated with each of the following clusters: ZG, ZF, ZX, Medulla, Fibroblast, White Adipose Tissue (WAT), Brown Adipose Tissue (BAT), Endothelial Cells, Erythrocytes, and White Blood Cells

## Author contributions

Conceptualization, M.B., and M.R.; Methodology, M.B., S.H., J.S.-Z., and M.R.; Investigation, M.B., S.H., M.S., A.Plewinski., and M.R.; Visualization, M.B, S.H., and M.R.; Resources, M.B., A.Pławski and M.R.; Writing—Original Draft Preparation, M.B., S.H., L.K.M., and M.R.; Writing—Review & Editing, all authors; Supervision and Funding Acquisition, L.K.M., A.Porzionato, and M.R.

## Institutional Review Board Statement

The animal study protocol was approved by the Local Ethics Committee for Animal Studies, Poznan, Poland (protocol No. 27/2023 from 17th April 2023).

## Code availability

Code used for data analysis has been deposited at GitHub (https://github.com/MarcinRuc78/Mouse-Adrenal-Spatial-Transcriptomic.git).

## Data availability

All datasets have been deposited in the Gene Expression Omnibus (GEO) database under accession number GSE283302.

https://adrenal-spatall-transcriptomic.shinyapps.io/Adrenal/

